# Memory encoding begins with spontaneous neural reactivation

**DOI:** 10.64898/2026.08.31.747822

**Authors:** Gemma Fernández-Rubio, Elvira Brattico, Morten L. Kringelbach, Peter Vuust, Leonardo Bonetti

**Affiliations:** Center for Music in the Brain, Department of Clinical Medicine, Aarhus University & The Royal Academy of Music, Aarhus, Aalborg, Denmark; Department of Education, Psychology, Communication, University of Bari Aldo Moro, Bari, Italy; Centre for Eudaimonia and Human Flourishing, Linacre College, University of Oxford, Oxford, United Kingdom; Department of Psychiatry, University of Oxford, Oxford, United Kingdom

**Keywords:** Long-term memory, Neural reactivation, Magnetoencephalography, Multivariate pattern analysis, Principal component analysis

## Abstract

Long-term memory relies on neural reactivation mechanisms that support consolidation and retrieval. However, it remains unknown whether these mechanisms also operate during encoding, the initial stage of memory formation. Using a decoding classifier, we analyzed magnetoencephalography (MEG) data from 67 participants as they actively encoded or passively listened to brief melodies. We found that the classifier generalized to the rest period immediately following encoding, revealing spontaneous neural reactivation and stable patterns after stimulus offset. This extends prior evidence of neural replay, which is typically observed during sleep and rest, by demonstrating reactivation during encoding, within the task itself. Decoding accuracy during encoding was correlated with recognition performance in subsequent tests, reinforcing its role in memory, with the largest differences between low- and high-recognition performance groups emerging 10 days after encoding. Whole-brain network analysis identified two primary systems: an auditory network (primary auditory cortex, insula, superior temporal gyrus) showing few condition differences, and a memory/associative network (hippocampus, parahippocampal gyrus, cingulate cortex) driving the neural reactivation underlying successful encoding. Together, these results demonstrate that spontaneous neural reactivation supports memory encoding even before formal consolidation.

## INTRODUCTION

One of the most remarkable features of human cognition is the ability to encode, store, and retrieve information, facilitating long-term memory. This capacity underlies not only everyday learning and decision-making but also the sophisticated mental processes that allow humans to adapt to and thrive in a complex environment ^1^. A thorough understanding of the neural mechanisms that enable memory formation, storage, and retrieval is thus critical for advancing knowledge of cognitive function and brain health.

At the systems level, memories are not simply stored in their final form at the time of encoding. Instead, additional processes, such as neural reactivation, play a pivotal role in transforming encoded information into long-lasting memories ^2,3^. Neural reactivation refers to the re-emergence of neural pathways that were initially engaged during learning ^4^. One specific form of reactivation is neural replay, which is the temporally structured, sequential reinstatement of neural patterns, often occurring at a faster pace and during sleep ^5–7^. Extensive animal work, particularly in rodents, has demonstrated that hippocampal replay during sleep supports the consolidation of spatial information ^8–10^. Neural replay has also been observed in the human motor cortex during sleep, indicating that replay is involved in reinforcing newly learned motor skills ^6,11,12^. Moreover, this phenomenon has been recorded during periods of quiet wakefulness, suggesting that consolidation processes extend beyond sleep ^13–16^.

Neural reactivation during memory retrieval has been well documented in human studies, reinforcing the idea that memory traces are dynamically reinstated ^17^. Retrieving a memory recruits the same brain regions engaged during encoding and enhances retention of information by reactivating both target and contextually linked content ^18,19^. Furthermore, the degree of neural reactivation has been correlated with measures of subjective vividness ^20^. In rodents, reactivation of cortical representations during memory retrieval can persist over weeks ^21^.

These findings have significantly expanded our understanding of the neural mechanisms underlying different memory processes, highlighting that the stages of consolidation and retrieval are dynamic periods during which memories are strengthened, reorganized, and integrated into existing knowledge networks ^2,7^. However, the immediate post-encoding period, before consolidation begins, remains underexplored. While neural replay during rest and sleep, as well as reactivation during retrieval, have been well characterized, it is unknown whether neural reactivation occurs in real time shortly after learning, and whether such spontaneous reactivation predicts long-term memory success.

To address this gap, we investigated whether neural reactivation emerges already at the stage of encoding. During magnetoencephalography (MEG) recordings, participants actively encoded or passively listened to brief musical sequences. By training a decoder to distinguish neural activity patterns between the two conditions, we were able to test for the presence of these patterns in the immediate post-encoding period. Critically, we hypothesized that spontaneous neural reactivation would be associated with better long-term recognition performance, providing novel evidence for a direct link between neural reactivation and memory outcomes.

## RESULTS

### Overview of experimental design and data analysis

Sixty-seven healthy participants underwent magnetoencephalography (MEG) and structural magnetic resonance imaging (MRI) scanning (**Figure 1a**). During MEG recordings, participants performed an auditory encoding task, where they were presented with the cue *listen* or *memorize* (1.5 s) before hearing a piano melody (2.5 s), followed by a period of silence (5.5 s) before the onset of the next trial (**Figure 1b**). Participants were instructed to passively listen to the melodies on *listen* trials and to actively memorize them on *memorize* trials. Six melodies, matched for musical features, were repeated 20 times, producing 60 *listen* trials and 60 *memorize* trials. Participants were randomly assigned to one of two groups, with each group listening and memorizing different sets of melodies (**Figure 1c**). Outside the MEG scanner, participants performed an old/new auditory recognition task similar to the one used in our previous studies ^22–25^ (**Figure 1d**). The task was administered on the same day as the MEG recordings, one day, and 10 days later.

**Figure 1.**
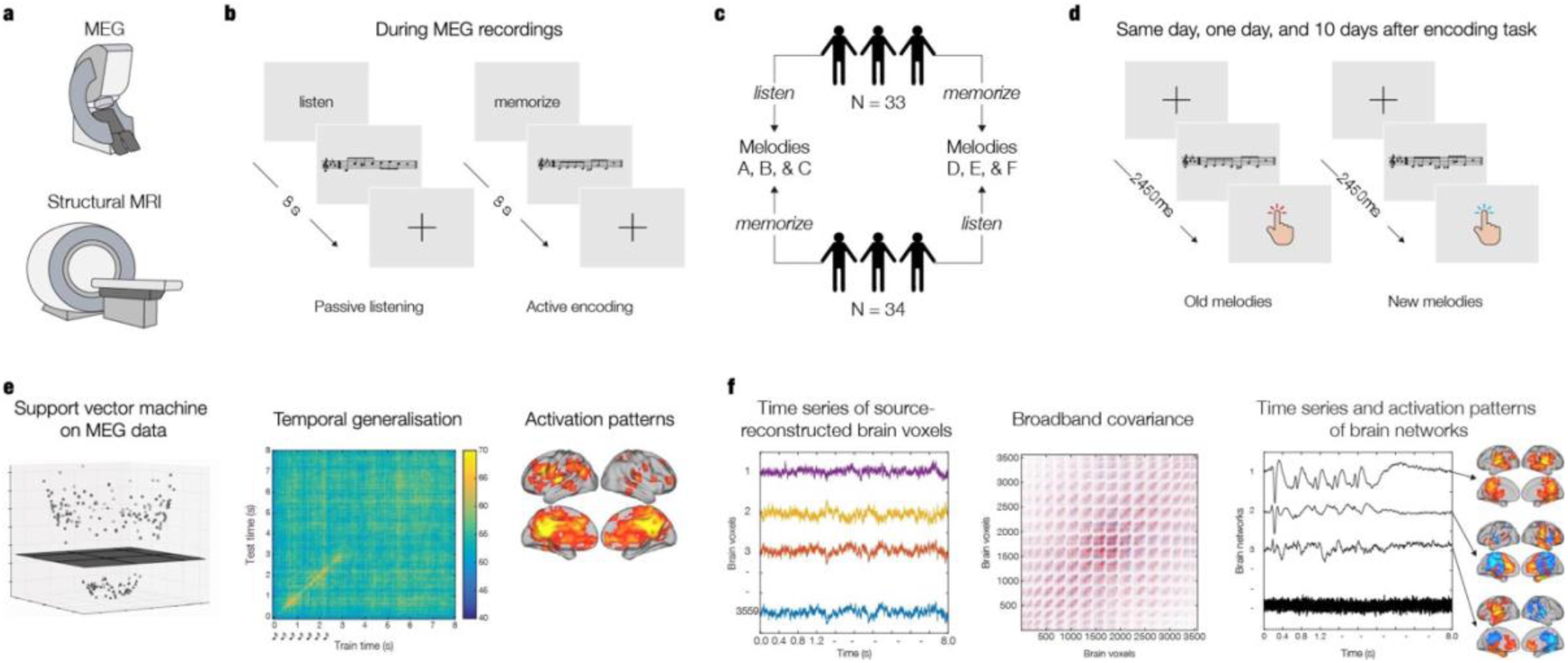
Experimental design and data analysis. **a) Data acquisition.** Magnetoencephalography (MEG) was used to collect neurophysiological data, while structural magnetic resonance imaging (MRI) was used to acquire anatomical brain scans. **b) Auditory encoding task.** An auditory encoding task was administered during MEG recordings. Six distinct melodies with matching musical features were presented, and participants were instructed to passively listen or actively memorize them. **c) Group counterbalancing.** Participants were randomly assigned to one of two conditions: one group passively listened to melodies A, B, and C, and actively memorized melodies D, E, and F, while the other group passively listened to melodies D, E, and F, and actively memorized melodies A, B, and C. **d) Old/new auditory recognition task.** Following the auditory encoding task, participants performed an old/new auditory recognition task outside the MEG. Participants were presented with the same melodies and with new melodies and were instructed to categorize them as old or new, respectively. The old/new auditory recognition task was performed on the same day, one day, and 10 days after the auditory encoding task. **e) Multivariate pattern analysis.** Multivariate pattern analysis was used to assess whether it was possible to discriminate the two experimental conditions (listen versus memorize) based on the neural activity recorded with MEG. The MEG data was co-registered with the MRI data, and source reconstruction was performed on the activation patterns emerged from the multivariate pattern analysis. **f) Whole-brain network analysis.** The BROADband brain Network Estimation via Source Separation (BROAD-NESS) pipeline was used to extract whole-brain networks from the source-reconstructed MEG data.

Multivariate pattern analysis was employed at single-participant level to decode the neural representations associated with passive listening and active encoding (**Figure 1e**). A support vector machine (SVM) classifier was used to calculate independent analyses for each participant ^22,26^. This analysis was expanded to a temporal generalization case to assess whether the experimental conditions were differentiated by neural patterns that were stable over time ^27,28^. The significance of the output was evaluated using a signed-rank permutation test against chance level (50%) for each timepoint, using false-discovery rate (FDR) correction for multiple comparisons. The multivariate pattern analysis produced decoding weights, which were later used to generate spatial activation patterns ^22,29^ (see Methods for details).

Further analyses were conducted to inspect the decoding accuracy in relation to the response accuracy data obtained from the old/new auditory recognition task. The sample was split into low- and high-performance groups based on the median response accuracy (i.e., number of correctly identified melodies) on each session of the task (same day, one day, and 10 days after encoding). Two-sample t-tests were computed to discriminate the decoding accuracy between the two groups independently for each session. Multiple comparisons were corrected for using cluster-based permutation tests ^30^.

The BROADband brain Network Estimation via Source Separation (BROAD-NESS) pipeline from the NESS framework ^31,32^ was employed to characterize whole-brain networks from the source-reconstructed MEG data (**Figure 1f**). After computing principal component analysis (PCA) and the time series of the brain networks for each participant and experimental condition, two-sided t-tests were computed for each time point and brain network to contrast the time series of the two conditions, using cluster-based permutation tests for correcting for multiple comparisons ^30^.

### Temporal and spatial differentiation of active encoding versus passive listening

The performance of the SVM classifier was significantly above chance level at 22.36% of timepoints (*p* < .011, FDR-corrected), indicating that the two conditions were discriminable on the basis of the neural activity generated. Differences extended beyond the diagonal of the temporal generalization matrix, indicating that distinct neural representations between active encoding and passive listening persisted after the melody ended, revealing spontaneous neural reactivation (**Figure 2a**).

**Figure 2.**
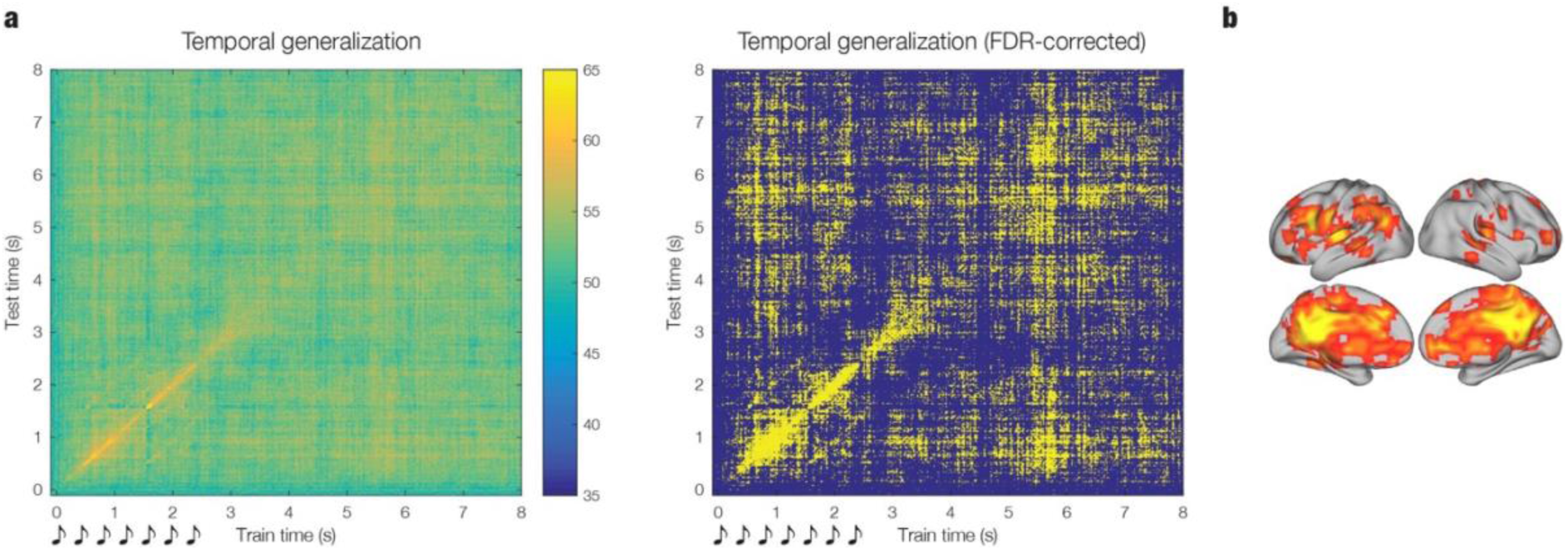
Neural activity associated with active encoding versus passive listening. **a) Decoding accuracy.** A temporal generalization matrix (tested against a 50% chance level) assessed whether the active encoding and passive listening conditions were differentiated by stable neural patterns (average of 67 participants). The color bar indicates decoding accuracy (%). Musical sketches below the matrix represent the onset of the notes comprising the melodies. The temporal generalization matrix was binarized after false-discovery rate (FDR) correction (p < .011). Significant timepoints are depicted in yellow. **b) Spatial activation patterns.** Main spatial activation patterns (i.e., brain sources larger than mean plus one standard deviation), contributing to the classification of the two experimental conditions (average of 67 participants).

The spatial activation patterns emerged from the decoding analysis signaled the brain regions that discriminated the two experimental conditions. These were primarily the bilateral cingulate gyrus and precuneus, as well as the left insula and left inferior frontal gyrus, which were consistently active throughout the trials (see **Table S1** for details). Main spatial activation patterns (i.e., brain sources larger than mean plus one standard deviation) were averaged across participants and over time (**Figure 2b**).

### Decoding accuracy as a function of long-term recognition performance

Decoding accuracy was examined in relation to performance on the old/new auditory recognition task. Participants were divided into low- and high-performance groups based on median response accuracy. The recognition task was administered on the same day as the encoding task, one day later, and 10 days later. For both groups and across all sessions, SVM classification performance exceeded chance level at multiple timepoints, both while the melodies were presented and in the period immediately after (see **Table S2** for details). Decoding accuracy was consistently higher in the high-performance group, particularly on the last test day, with significant group differences at 0.17% of timepoints on the same day as encoding (**Figure 3a**), 0.62% of timepoints one day later (**Figure 3b**), and 5.70% of timepoints 10 days later (**Figure 3b**).

**Figure 3.**
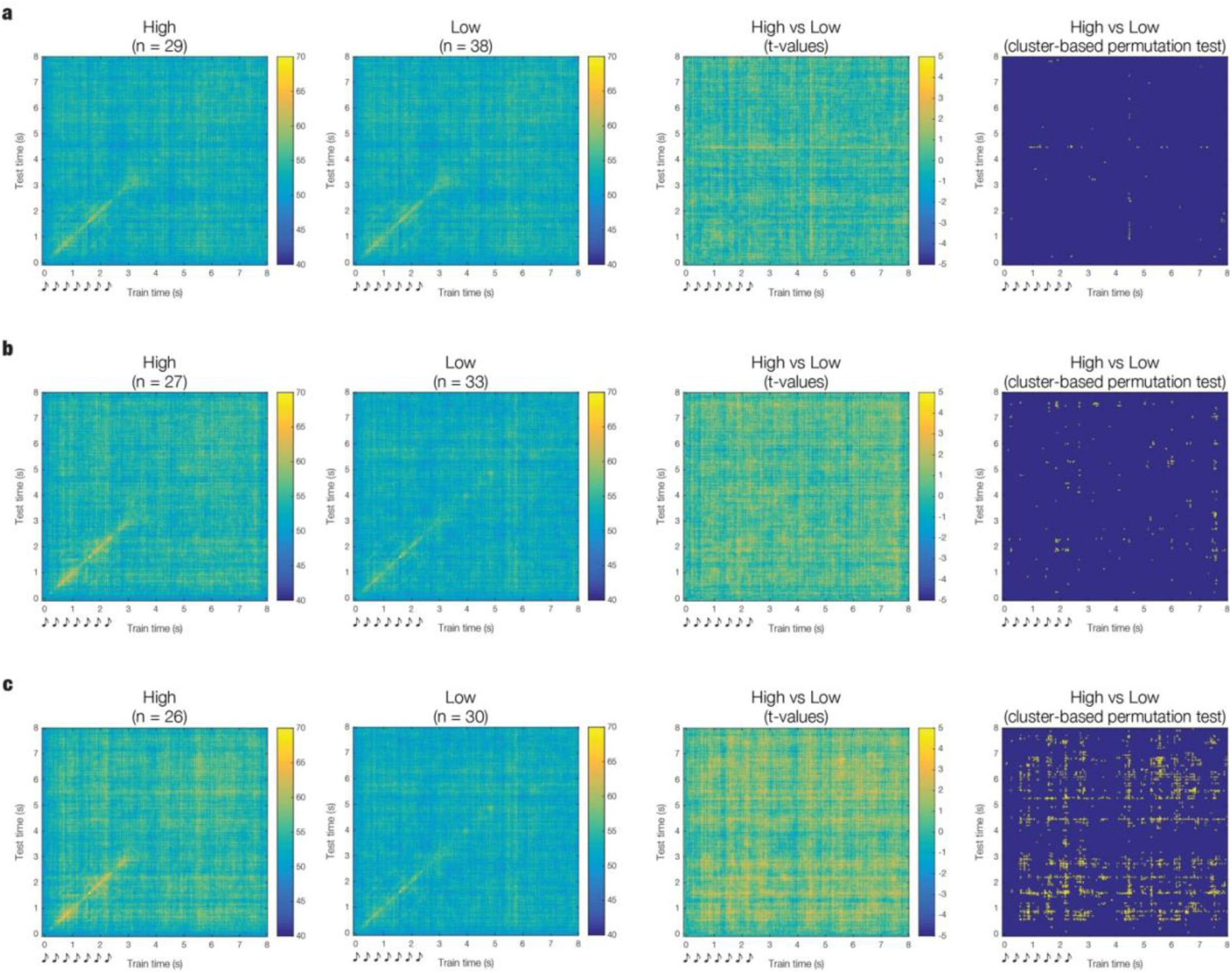
Differences in decoding accuracy between low- and high-performance recognition groups. Temporal generalization matrices (tested against a 50% chance level) assessed whether the active encoding and passive listening conditions were differentiated by stable neural patterns. These were computed independently for each performance group based on median response accuracy data from the old/new auditory recognition task. Color bars indicate decoding accuracy (%). Group differences in decoding accuracy were estimated using two-sided t-tests, correcting for multiple comparisons with cluster-based permutation tests. Musical sketches below matrices represent the onset of each note comprising the melodies. **a) Same day as encoding. b) One day after encoding. c) Ten days after encoding.**

### Functional brain networks during active encoding and passive listening

Whole-brain networks were estimated using PCA within the BROAD-NESS framework. The effective dimensionality was 2.52, revealing two principal components (i.e., brain networks) accounting for 60.55% and 16.32% of the variance, with a third component approaching the threshold at 4.74% (see **Methods** and **Figure 4a**).

**Figure 4.**
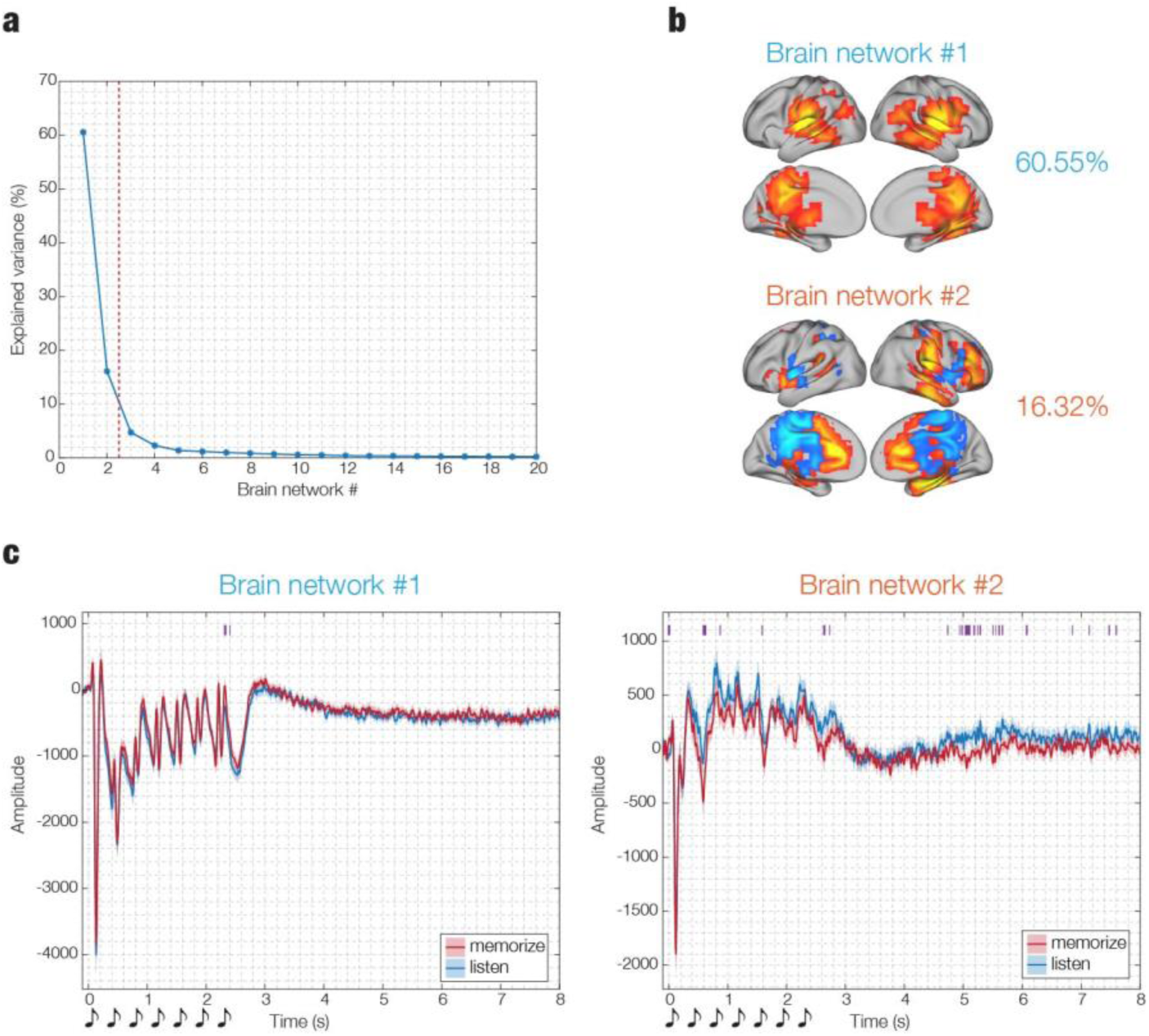
Key brain networks associated with active encoding and passive listening. **a. Explained variance.** Whole-brain networks were estimated via principal component analysis (PCA) within the BROADband brain Network Estimation via Source Separation (BROAD-NESS) pipeline. The red dotted line marks the effective dimensionality. **b. Spatial activation patterns.** Brain networks with the highest explained variance (60.55% and 16.32%, respectively). Brain templates illustrate the spatial extent of the networks. Yellow voxels contribute most, and light blue voxels contribute least, to the time series. **c. Time series.** Time series of the two brain networks that explained the highest variance, estimated independently for each participant and experimental condition. Individual time series were averaged across participants (n = 67). Shaded areas represent standard errors. Purple lines above the plots mark the temporal extent of significant differences between conditions.

The PCA-derived spatial activation patterns revealed that the first brain network primarily comprised auditory regions, including the primary auditory cortices, insulae, and superior temporal gyri. The second brain network comprised mainly memory regions (hippocampus and parahippocampal gyri) and associative regions (medial and anterior cingulate gyri) (**Figure 4b** and **Table S4**). The third brain network displayed widespread activation in both auditory and memory regions (**Figure S1a** and **Table S4**).

Independent time series were generated for each participant, brain network, and experimental condition using PCA-derived weights from the averaged data. Two-sided t-tests were computed to contrast the time series of the two conditions independently for each timepoint and brain network, using cluster-based permutation tests for correcting for multiple comparisons (1000 permutations, *α* = 0.05). The second brain network showed the most significant differences between conditions, both during and immediately after melody presentation, indicating its relevance for the task. Fewer significant differences were observed in the first and third networks (**Figure 4c**, **Figure S1a**, and **Table S3**).

### Differences in long-term recognition across sessions

During the old/new auditory recognition task, participants classified melodies as old or new. To account for differences between conditions, old melodies were later subdivided into actively memorized (*old*) and passively listened (*listen*) conditions. Response accuracy data from this task were then analyzed across three conditions (*old*, *listen*, *new*) and three sessions (same day, one day, 10 days after encoding). A binomial generalized linear mixed model (GLMM) with fixed effects for condition, time, and their interaction, plus random subject intercepts, was used. Post-hoc pairwise comparisons were performed using Tukey adjustments.

The GLMM revealed a significant time and condition × time interaction effects , while the condition effect was non-significant. Post-hoc comparisons showed that success rates were higher at session 1 compared to session 2 (*p* = .005) and session 3 (*p* < .001), with no difference between sessions 2 and 3 (*p* = .689). The significant interaction indicated that differences between conditions varied over time. At session 1, no differences between conditions were observed. At session 2, success rates were higher for *new* > *old* ( *p* < .001), *new* > *listen* (*p* < .001), and *old* > *listen* (*p* < .044). At session 3, success rates were higher for *new* > *listen* (*p* < .001) and marginal differences were observed for *old* > *listen* (*p* = .052) (**Figure S2**).

## DISCUSSION

The goal of this study was to provide new insights into the neural mechanisms of memory formation by exploring whether spontaneous neural reactivation takes place immediately after encoding. The decoder trained to differentiate between neural patterns during active encoding versus passive listening of the melodies was able to generalize its predictions to the few seconds immediately following the presentation of the stimuli. During this period, participants were simply instructed to rest, awaiting the next trial. This finding is significant because it reveals that the neural patterns associated with encoding remain stable even after the stimuli have been removed, suggesting the presence of a sustained neural process during this brief rest period. Importantly, the classifier distinguished between active encoding and passive listening, despite the melodies being matched on all musical features, providing compelling evidence that the neural patterns reflect higher-level cognitive processes associated with active engagement and memory formation. This result effectively rules out auditory perception as the primary driver, underscoring the critical role of cognitive involvement in memory encoding.

These findings align with the extensive literature on neural reactivation. In rodents, neural replay is a well-documented phenomenon, particularly in the hippocampus, where neurons that were activated during spatial navigation tasks are sequentially reactivated during rest and sleep ^8–10^. This mechanism has been shown to be crucial for the consolidation of memories, facilitating long-term retention and recall ^2,3^. In humans, studies demonstrated that neural replay takes place during sleep and quiet wakefulness, contributing to the consolidation of memories, particularly in tasks involving motor learning ^11,12^. Furthermore, research applying multivariate pattern analysis to MEG recordings during non-spatial reasoning tasks demonstrated the occurrence of spontaneous visual representations in fast sequences during rest periods ^33–35^. Beyond memory consolidation, neural reactivation during memory retrieval enhances information retention ^18,19^. Our findings expand upon this body of work by demonstrating that neural reactivation can occur spontaneously already at the initial learning stage. Specifically, we observed neural reactivation during the short intervals immediately following the encoding phase, when participants waited for the next sequence to be presented. This suggests that spontaneous neural reactivation may play a role in memory encoding, even before any formal consolidation period begins. Future studies should investigate whether the reactivation reflects a general phenomenon or if it involves the replay of specific musical sequences in the same temporal order as they were initially presented. This would provide valuable insights into the precision of neural replay during encoding.

Analysis of the neural sources that contributed to the decoding revealed significant involvement of the bilateral cingulate gyrus and precuneus. These regions have been previously implicated in music processing and are known to play roles in integrating sensory and cognitive processes, particularly in relation to attention and memory ^22,25,36–39^. The relevance of these regions in our study underscores their potential role in encoding complex auditory information, such as musical sequences. Other areas that contributed to the decoding were the left insula and left inferior frontal operculum, in line with our previous research on the encoding and recognition of musical tones ^25,40–43^. Additionally, there was some involvement of the left hippocampus and left parahippocampal gyrus, which are structures in the medial temporal lobe that are key for memory ^44,45^.

The results from the whole-brain analysis within the BROAD-NESS framework complement and expand the findings on the spatial activation patterns from the multivariate pattern analysis. Similar to our previous study on memory recognition ^31^, we identified two primary networks: one comprising auditory regions (primary auditory cortex, insula, superior temporal gyrus), and another encompassing memory and associative regions (hippocampus, parahippocampal gyrus, medial cingulate gyrus). While the majority of differences between conditions emerged in the latter brain network, we observed striking similarities between the two networks (which together explained 76.87% of the variance) and the spatial activation patterns derived from the decoding. Both analytical approaches converge on a consistent picture: memory and associative regions, particularly the cingulate gyrus, precuneus, hippocampus, and parahippocampal gyrus, play a central role in encoding musical sequences, while auditory regions, particularly the insula and primary auditory cortex, are crucial to both encoding and passive listening. The decoding captures regions where the two conditions differ across the full time window, while the whole-brain analyses provide a mechanistic explanation for how these brain regions coordinate into networks that drive the spontaneous neural reactivation underlying successful encoding.

Finally, a critical finding was that decoding accuracy during encoding predicted subsequent recognition performance. This effect increased across sessions, with the largest difference between low- and high-performance groups occurring 10 days after encoding. These results reinforce the idea that the neural patterns identified by the classifier are indeed reflective of encoding processes and are predictive of long-term memory success, consistent with the role of neural replay and neural reactivation in strengthening representations during memory consolidation and retrieval, respectively ^17,46^. The behavioral results mirrored decoding accuracy in that both declined over time, peaking in the first session. Moreover, recognition accuracy was highest for *old* and *new* responses compared to *listen* responses, demonstrating that the classifier captured behaviorally relevant neural signatures of memory formation.

To our knowledge, our findings provide the first evidence that neural reactivation occurs immediately after encoding and predicts enhanced long-term memory recognition, establishing a novel mechanism in memory formation. Future studies should contrast this spontaneous reactivation with neural replay during extended rest and investigate how both processes contribute to neural mechanisms underlying later retrieval, as we have shown during recognition of musical sequences ^22–25,47^. By addressing these questions, future work could significantly advance our understanding of how the brain encodes, consolidates, and retrieves memories.

## METHODS

### Sample

Seventy-one healthy volunteers were recruited for the study. From this sample, four were excluded due to noisy or corrupted neural data, yielding a final sample of 67 participants (47 female, mean age = 25.88 ± 6.91). Participants were recruited in Denmark, primarily from Western countries, and reported similar educational backgrounds: 22 participants held a bachelor’s degree, 22 held a master’s degree, 16 held a high school diploma, 5 held a professional degree, and two did not complete high school.

Participants gave their explicit informed consent and received a small monetary compensation. The study was approved by the Institutional Review Board (IRB) of Aarhus University (case number: DNC-IRB-2022-011) and conducted in accordance with the Declaration of Helsinki - Ethical Principles for Medical Research.

### Experimental stimuli and design

Neurophysiological data were collected with magnetoencephalography (MEG) and structural brain scans were acquired using magnetic resonance imaging (MRI) (**Figure 1a**). An auditory encoding task was administered during MEG recordings. On each trial, participants were presented with the words *listen* or *memorize* on the screen for 1.5 seconds and subsequently heard a melody that lasted 2.5 seconds and was paired with a fixation cross on the screen. This was followed by a period of 5.5 seconds of silence before the onset of the next trial. There were 120 trials (60 listen and 60 memorize) presented in random order, and each melody was repeated 20 times (**Figure 1b**).

Six different melodies were used in the experiment. The melodies were created by GFR and matched in terms of the duration, rhythm, volume, timbre, tempo, and meter. Each melody consisted of seven notes played on a piano, with the first note always being middle C (261 Hz). An isochronous rhythmic pattern (i.e., the time intervals between notes are equally spaced) was used and each note lasted 350 milliseconds. MIDI versions of the melodies were created on MuseScore v3.6.2 and presented using Psychopy v3.0. Participants were randomly assigned to one of two groups, with each group listening and memorizing different sets of melodies. Group 1 (n = 33) passively listened to melodies A, B, and C, and actively memorized melodies D, E, and F, while Group 2 (n = 34) did the opposite (**Figure 1c**).

Outside the MEG scanner, participants performed an old/new auditory recognition task ^22–25,43,48–50^ (**Figure 1d**). Participants were presented with the same melodies from the encoding task (i.e., old melodies) and with variations of these (i.e., new melodies). For each of them, they judged whether the melody was old or new by pressing one of two keys. Six original melodies were used (repeated twice) and 12 variations were created. Participants performed this task on the same day as the MEG recordings, one day later, and 10 days later. For logistic limitations, MEG data was not acquired during the recognition task. The task was performed at Aarhus University Hospital on the first day, and online on the second and tenth days using the software Cognition (www.cognition.run).

### Data acquisition

MEG data were recorded in a magnetically shielded room at Aarhus University Hospital (AUH), Aarhus, Denmark, using a 306-channel TRIUX MEG scanner (Elekta Neuromag, Helsinki, Finland). Data were acquired at a sampling rate of 1000 Hz with an analogue filtering of 0.1–330 Hz. The participants’ head shape and position of four Head Position Indicator (HPI) coils were registered with a 3D digitizer (Polhemus Fastrak, Colchester, VT, USA) before the MEG scanning. The HPI coils continuously registered the position of the head during MEG recordings, which was later used for movement correction. Finally, two sets of bipolar electrodes were used to record eye movements and cardiac rhythm, which were later used to remove electrooculography (EOG) and electrocardiography (ECG) artefacts.

MRI data were recorded on a CE-approved 3T Siemens MRI-scanner at AUH. Structural T1 (mprage with fat saturation) with a spatial resolution of 1.0 × 1.0 × 1.0 mm were acquired using the following sequence parameters: echo time (TE) = 2.61 ms, repetition time (TR) = 2300 ms, reconstructed matrix size = 256 × 256, echo spacing = 7.6 ms, bandwidth = 290 Hz/Px.

### MEG data preprocessing

Raw MEG sensor data (204 planar gradiometers and 102 magnetometers) was first preprocessed by MaxFilter version 2.2.15 ^51^ to attenuate external interferences. The following MaxFilter parameters were applied: spatiotemporal signal space separation (SSS), down-sample from 1000 Hz to 250 Hz, movement compensation using HPI coils (default step size: 10 ms), and a correlation limit of 0.98 to reject overlapping signals between inner and outer subspaces during spatiotemporal SSS.

Data were subsequently converted into Statistical Parametric Mapping (SPM) format and further preprocessed and analyzed in MATLAB (MathWorks, Natick, MA, USA) using a combination of in-house-built codes (LBPD, https://github.com/leonardob92/LBPD-1.0.git) and the Oxford Centre for Human Brain Activity (OHBA) Software Library OSL; ^52^, which utilizes Fieldtrip ^53^, FSL^54^, and SPM12 ^55^ toolboxes. Continuous MEG data were visually inspected to remove artifacts using the OSLview tool. Independent component analyses (ICA) were used to discard the interference of eye movements and cardiac rhythm from the brain data (OSL implementation).

A 30 Hz low-pass filter was applied to the continuous data to remove high-frequency noise, in accordance with standard decoding practices ^56,57^. Finally, the signal was epoched in 120 trials (60 listen, 60 memorize) and baseline-corrected by removing the mean signal recorded in the baseline from the post-stimulus brain signal. Each trial lasted 8.1 seconds (100 ms of baseline plus 2450 ms of auditory stimulation plus 5450 ms of silence).

### Multivariate pattern analysis

Multivariate pattern analysis was employed to classify (i.e., decode) the neural representations associated with passive listening and active encoding (**Figure 1e**). A support vector machine (SVM) classifier was used to calculate independent analyses for each participant ^22,26^. MEG sensor data were rearranged into a 3D matrix (306 channels x 2026 timepoints x 120 trials) and submitted to the SVM classifier to discriminate between the two experimental conditions. A leave-one-out cross-validation approach was adopted to avoid overfitting. This procedure divided the trials into five different groups and assigned, for each timepoint, four groups to the training set and the remaining single group to the testing set. This process was performed 100 times with random reassignment of the data to training and testing sets. The decoding accuracy time series obtained for each permutation was then averaged to obtain a final time series showing the performance of the classifier for each participant.

To assess whether the experimental conditions were differentiated by neural patterns that were stable over time, the decoding analysis reported above was expanded to a temporal generalization case ^27,28^. Here, the SVM algorithm used each timepoint of the training set to predict all timepoints in the testing set. The significance of the output was evaluated using a signed-rank permutation test against chance level (50%) for each timepoint (α = .05). Multiple comparisons were corrected using false-discovery rate (FDR) correction (FDR-adjusted *p* < .011).

### Source reconstruction and spatial activation patterns

The multivariate pattern analysis returned decoding weights independently for each timepoint. First, the spatial activation patterns at MEG sensor level were computed by multiplying the covariance matrix of the sensor data by the decoding weights ^29^. Second, the sources of the recorded MEG signal were reconstructed. Using the information collected with the 3D digitizer, the MEG data and individual MRI T1-weighted images were co-registered, and the forward model was subsequently computed. A single-shell forward model using an 8-mm grid and beamforming algorithm as inverse solution were employed ^58–60^. The weights obtained from the beamforming algorithm were then multiplied by the spatial activation patterns obtained from the decoding analysis to return their spatial projection in source space, independently for each timepoint ^29^. This procedure returned a 2D matrix (3559 brain sources x 2026 timepoints). For illustrative purposes, the spatial activation patterns were averaged over time and participants and depicted in brain templates.

### Multivariate pattern analysis by groups

Further analyses were conducted to inspect the decoding accuracy in relation to the response accuracy data obtained from the old/new auditory recognition task. The sample was split into low- and high-performance groups based on the median response accuracy (i.e., number of correctly identified melodies). This resulted in 38 participants in the low-performance group and 29 participants in the high-performance group on the first session (same day as the encoding task), 33 participants in the low-performance group and 27 participants in the high-performance group on the second session (one day after the encoding task), and 30 participants in the low-performance group and 26 participants in the high-performance group on the third session (10 days after the encoding task). To be noted, seven participants did not complete the old/new auditory recognition task on the second session (n = 60) and 11 participants dropped out on the third session (n = 56).

For each session, differences between the low- and high-performance groups were tested based on the outcome of the multivariate pattern analysis computed earlier. To this aim, two-sample t-tests were computed to discriminate the decoding accuracy between the two groups independently for each session. Multiple comparisons were corrected for using cluster-based permutation tests ^30^. First, a two-sample t-test was computed independently for each timepoint of the temporal generalization matrices. Clusters of neighboring significant timepoints were then identifyed by applying a threshold to the statistical results (*α* = 0.05). Next, 100 permutations were performed, where, for each permutation, the labels of the two groups were shuffled for each participant, and the statistics were recalculated. A reference distribution of significant clusters was built and compared with the clusters from the original data. Clusters in the original data were deemed significant only if their sizes exceeded the maximum cluster size observed in 95% of the permuted data.

### Whole-brain network analysis

The BROADband brain Network Estimation via Source Separation (BROAD-NESS) pipeline from the NESS framework ^31,32^ was employed to characterize whole-brain networks from the source-reconstructed MEG data (**Figure 1f**). The time series data from the 3559 source-reconstructed brain voxels was used to calculate the covariance matrix (C). Principal component analysis (PCA) was then computed by solving the eigenequation CW = WΛ for the eigenvectors (W) to find the weighted combinations of brain voxels to orthogonal components, which represent brain networks, while the associated eigenvalues (Λ) express the amount of variance explained by each network component. Network activation time series (y) were derived by applying the spatial filters (W) to the voxel data matrix (X). The same filters were applied to the covariance matrix (C) to compute the spatial activation patterns (for details on this procedure, see ^31^).

After computing PCA and the time series of the brain networks for each participant and experimental condition, two-sided t-tests were computed for each time point and brain network to contrast the time series of the two conditions (*α* = 0.05). Multiple comparisons were corrected with cluster-based permutation tests (1000 permutations) analogues to the ones described above ^30^.

### Behavioral data analysis

Behavioral data from the old/new auditory recognition task were analyzed to determine differences in response accuracy across experimental conditions and sessions. During the task, participants classified melodies as old or new. To account for differences between conditions, old melodies were later subdivided into actively memorized (old) and passively heard (listen) conditions. Thus, three conditions were analyzed: *old*, *listen*, and *new*.

A generalized linear mixed model (GLMM) with a binomial distribution and logit link was used to analyze success counts across the three experimental conditions, measured at three sessions (same day, one day, and 10 days after the encoding task). The model included fixed effects for condition, time, and their interaction, with random intercepts for subjects to account for repeated measures. Post-hoc pairwise comparisons were performed using Tukey adjustments.

## ACKNOWLEDGEMENTS

The Center for Music in the Brain (MIB) is funded by the Danish National Research Foundation (project number DNRF117), The Lundbeck Foundation (R469-2024-1573), and Købmand Herman Sallings Fond. L.B. is supported by Sapere Aude: Independent Research Fund Denmark (DFF) Research Leader (grant ID: 10.46540/5253-00003B), Lundbeck Foundation (Talent Prize 2022), Carlsberg Foundation (CF20-0239), Center for Music in the Brain, Linacre College of the University of Oxford, and Nordic Mensa Fund. M.L.K is supported by Center for Music in the Brain and Centre for Eudaimonia and Human Flourishing, which is funded by the Pettit and Carlsberg Foundations. We thank Nikita Joe, Shiraz Ben-Shoshan, and Samuel William Nehrer for their assistance with data collection.

## AUTHOR CONTRIBUTIONS

G.F.-R., E.B., and L.B. designed the study. L.B., M.L.K., and P.V. recruited the resources for the experiment. G.F.-R. collected the data. G.F.-R. and L.B. performed pre-processing, multivariate pattern analysis, and whole-brain network analysis. G.F.-R. and L.B. prepared the figures and wrote the first draft of the manuscript. All the authors contributed to and approved the final version of the manuscript.

## CODE AVAILABILITY

The main data analysis pipeline used in this study is available at the following link: https://github.com/gemmaferu/neural-reactivation

The BROAD-NESS toolbox is available at the following link: https://github.com/leonardob92/BROADNESS_MEG_AuditoryRecognition/tree/main/BROADNESS_Toolbox

## SUPPORTING INFORMATION

### Supplementary figures

**Figure S1.**
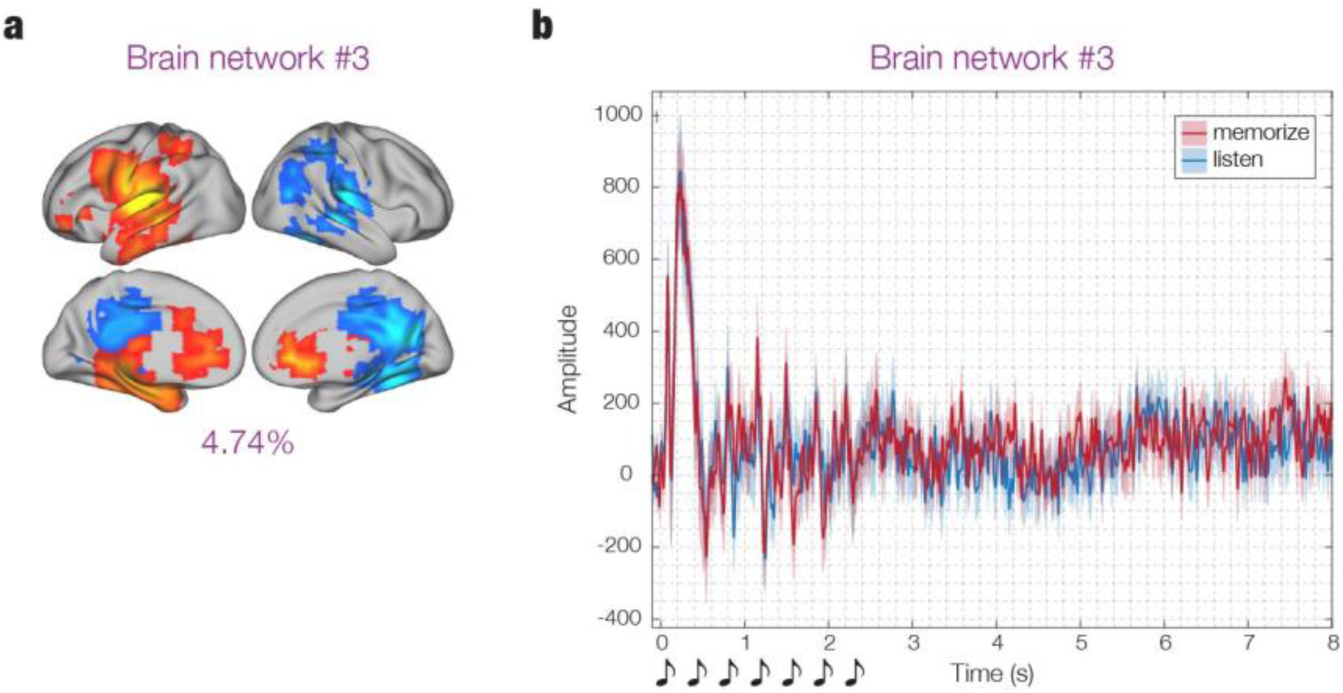
Brain network #3 associated with active encoding and passive listening. **a. Spatial activation patterns.** Brain network with the third highest explained variance (4.74%). Brain templates illustrate the spatial extent of the brain network. Yellow voxels contribute most, and light blue voxels contribute least, to the time series. **b. Time series.** Time series of the third brain network, estimated independently for each participant and experimental condition. The individual time series was averaged across participants (n = 67). Shaded areas represent standard errors. Purple lines above the plot mark the temporal extent of significant differences between conditions.

**Figure S2.**
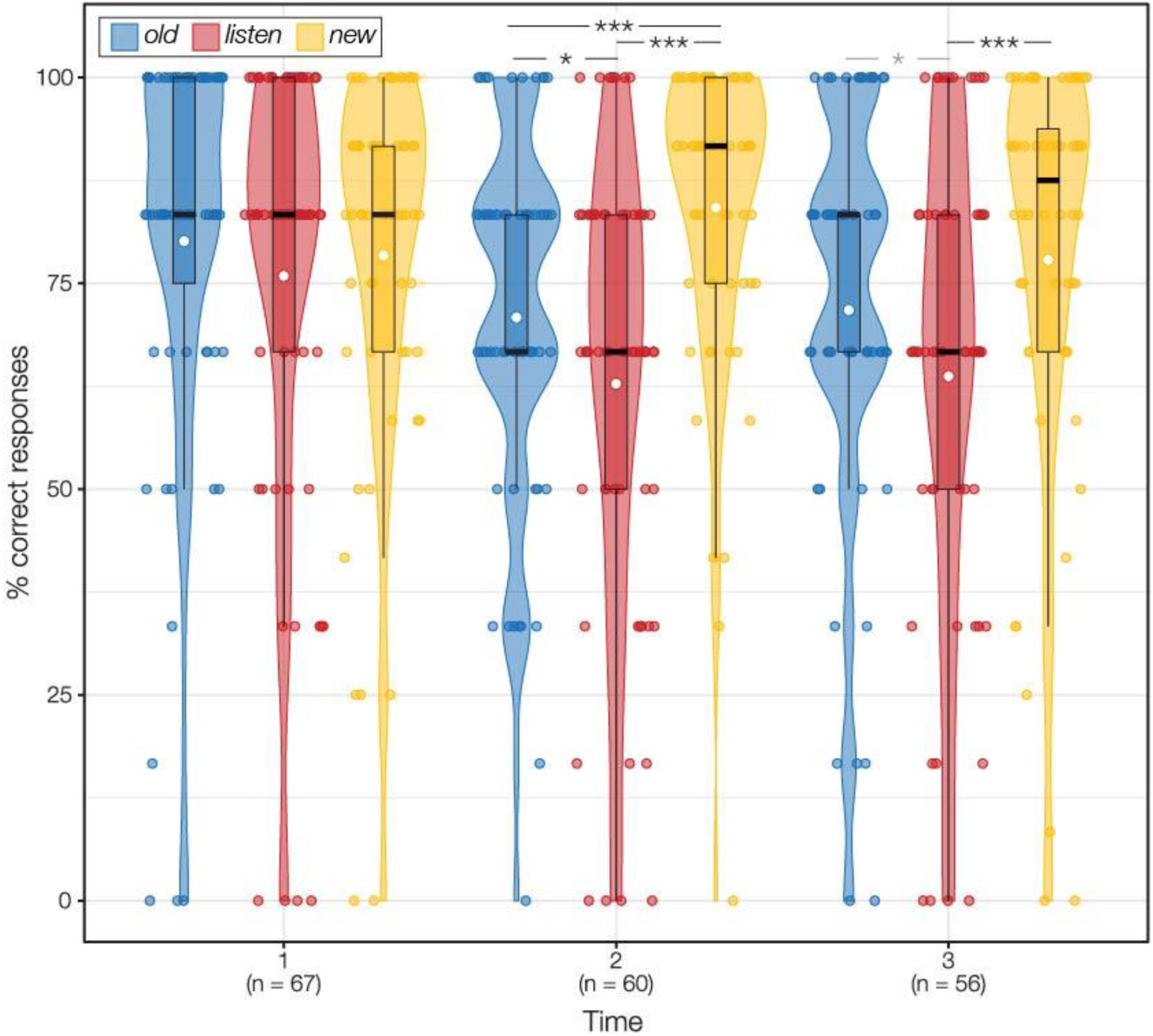
Response accuracy in old/new auditory recognition task. Violin plots with overlaid boxplots and jittered individual data points illustrating the distribution of correct response proportions in the old/new auditory recognition task across three experimental conditions (old, listen, new) measured at three timepoints (1 = same day as encoding task; 2 = one day later; 3 = 10 days later). White dots represent the mean proportion of correct responses for each condition and timepoint. Asterisks above the plots denote statistically significant differences from post-hoc tests of the interaction effect; a lighter-colored asterisk indicates a marginally significant difference (p = 0.052).

## Notes

### Competing Interest Statement

The authors have declared no competing interest.

